# *Etv5* is not required for Schwann cell development but is required to regulate the Schwann cell response to peripheral nerve injury

**DOI:** 10.1101/2020.09.23.309815

**Authors:** Anjali Balakrishnan, Lauren Belfiore, Lakshmy Vasan, Yacine Touahri, Morgan Stykel, Taylor Fleming, Rajiv Midha, Jeff Biernaskie, Carol Schuurmans

## Abstract

Schwann cells are the principal glial cells of the peripheral nervous system, and their development into myelinating glia is critically dependent on MEK/ERK signaling. Ets-domain transcription factors (*Etv1, Etv4, Etv5*) are common downstream effectors of MEK/ERK signalling, but so far, only *Etv1* has been ascribed a role in Schwann cell development, and only in non-myelinating cells. Here, we examined the role of *Etv5*, which is expressed in Schwann cell precursors, including neural crest cells and satellite glia, in Schwann cell lineage development. We analysed *Etv5*^*tm1Kmm*^ mutants (designated *Etv5*^*−/−*^) at embryonic days (E) 12.5, E15.5 and E18.5, focusing on dorsal root ganglia. At these embryonic stages, satellite glia (glutamine synthetase) and Schwann cell markers, including transcriptional regulators (Sox10, Sox9, Tfap2a, Pou3f1) and non-transcription factors (Ngfr, BFABP, GFAP), were expressed in the DRG of wild-type and *Etv5*^*−/−*^ embryos. Furthermore, by E18.5, quantification of Sox10^+^ Schwann cells and NeuN^+^ neurons revealed that these cells were present in normal numbers in the *Etv5*^*−/−*^ dorsal root ganglia. We next performed peripheral nerve injuries at postnatal day 21, revealing that *Etv5*^−/−^ mice had an enhanced injury response, generating more Sox10^+^ Schwann cells compared to wild-type animals at five days post-injury. Thus, while *Etv5* is not required for Schwann cell development, possibly due to genetic redundancy with *Etv1* and/or *Etv4, Etv5* is an essential negative regulator of the peripheral nerve injury repair response.

**SIGNIFICANCE STATEMENT:** Our study sought to determine whether the ets domain transcription factor, *Etv5*, plays a role in regulating Schwann cell development and nerve repair. By using an embryonically and postnatally viable hypomorphic *Etv5* mutant allele, we demonstrated that *Etv5* is not required for the development of Schwann cells or other neural crest derivatives in the dorsal root ganglia, including satellite glia and neurons. Surprisingly, loss of *Etv5* had a direct impact on the Schwann cell repair response post-injury, resulting in more Schwann cells populating the distal injured nerve site compared to wild-type animals. Thus, this work describes for the first time a role for *Etv5* in regulating the Schwann cell repair response after peripheral nerve injury.

## INTRODUCTION

Macroglial cells in the peripheral nervous system (PNS) include Schwann cells and satellite glial cells, which populate peripheral ganglia, including the dorsal root ganglia (DRG) that flank the developing spinal cord. Schwann cells myelinate peripheral nerves, but also wrap non-myelinated axons to regulate neuronal survival and axonal diameter (Jessen and Mirsky, 2005). Conversely, satellite glia encircle peripheral neuronal cell bodies, and have been likened to central nervous system (CNS) astrocytes (Kastriti and Adameyko, 2017). There is also growing support for a role for satellite glia in regulating pain (Jasmin et al., 2010; Ji et al., 2013; Song et al., 2018; Wang et al., 2019; Zhang et al., 2020). Despite their distinct positioning, Schwann cells (wrapping axons) and satellite glia (wrapping neuronal cell bodies) share an embryonic origin, both arising from neural crest cell (NCC) progenitors (Jessen and Mirsky, 2019a). Moreover, satellite glia can transition to a Schwann cell fate, at least *in vitro* (Cameron-Curry et al., 1993; Hagedorn et al., 2000), highlighting their close lineage relationship, and suggesting that satellite glia could replenish Schwann cells when required for repair.

Multiple signaling molecules and transcriptional regulators regulate Schwann cell development (Castelnovo et al., 2017). The ErbB family of receptor tyrosine kinases (RTKs), which are activated by Neuregulin 1 (NRG1), an EGF family ligand, have emerged as central regulators of Schwann cell precursor (SCP) proliferation, migration and myelination (Newbern and Birchmeier, 2010). In addition, the MEK-ERK signal transduction cascade, activated downstream of RTK signaling, is essential for Schwann cell differentiation, as revealed by the analysis of genetic mutants of ERK1 and ERK2 (Newbern et al., 2011). Conversely, if ERK signaling is ectopically activated in Schwann cells, myelination ensues (Ishii et al., 2013; Sheean et al., 2014).

ERK kinases have several downstream effectors that they regulate by phosphorylation, including multiple transcription factors (Tsang and Dawid, 2004), such as those of the Ets-domain (helix-turn-helix super family) family (Yang et al., 2013). In Drosophila, *Pointed* (*Pnt*) is an ets-domain transcriptional activator that is activated downstream of RTK signaling and which is required for glial cell fate specification (Klaes et al., 1994). In vertebrates, critical ets domain factors activated downstream of RTK signaling include *Etv1* (*ER81*), *Etv4* (*Pea3*) and *Etv5* (*Erm*). Activation of MEK-ERK signaling initiates the expression of *Etv1* and *Etv5* to specify an oligodendrocyte fate, which are myelinating glial cells in the CNS (Li et al., 2012; Wang et al., 2012; Li et al., 2014; Ahmad et al., 2019). *Etv1* is also expressed in Schwann cells (Srinivasan et al., 2007), but it is not required for the development of myelinating Schwann cells (Fleming et al., 2016). Instead, *Etv1* facilitates interactions between peripheral axons and non-myelinating Schwann cells in Pacinian corpuscles (Sedy et al., 2006; Fleming et al., 2016).

*Etv5* is also expressed in the developing PNS, beginning at embryonic day (E) 9.0 in NCCs and persisting until ∼ E12.5 in SCPs and in satellite glial cells (Hagedorn et al., 2000; Balakrishnan et al., 2016). *Etv5* expression is not glial specific, as it is also expressed in TrkA^+^ sensory neurons in peripheral ganglia (Chotteau-Lelievre et al., 1997; Hagedorn et al., 2000). Blocking *Etv5* function with a dominant negative construct in NCCs *in vitro* blocks neuronal fate specification but does not affect glial differentiation, NCC survival or proliferation (Paratore et al., 2002). However, whether *Etv5* is required for Schwann cell development *in vivo* has not been addressed.

Strikingly, several genes expressed during Schwann cells development are also expressed post peripheral nerve injury and support a proliferative repair Schwann cell phenotype (Balakrishnan et al., 2016). Here, we analysed *Etv5*^*tm1Kmm*^ mutant mice carrying a deletion of exons 2-5 (hereafter designated *Etv5*^−/−^) (Chen et al., 2005) to ask whether *Etv5* is required for Schwann cell development and the peripheral nerve injury response. We found no evidence for Schwann cell developmental defects in *Etv5*^−/−^ embryos. However, we did observe an expansion of Sox10^+^ Schwann cells in *Etv5*^−/−^ peripheral nerves post-injury. Thus, *Etv5* acts as a negative regulator of the Schwann cell repair response, and in its absence, more Schwann cells are generated. We discuss our findings in the context of important caveats, such as the potential for genetic redundancy with *Etv1* and/or *Etv4*, and our use of a hypomorphic mutant allele, which was necessitated by the early embryonic lethality of *Etv5* null mice.

## MATERIAL AND METHODS

### Animals and genotyping

*Etv5*^*tm1Kmm*^/J mice (Stock No. 022300) from Jackson Laboratory (ME, United States) were maintained on a 129/SvJ background. Mice were maintained as heterozygotes in a 12 hr light / 12 hr dark cycle. Heterozygous intercrosses were set up to generate homozygous mutant embryos, designated *Etv5*^*−/−*^. Pregnancy was determined by detection of a vaginal plug, with the morning of plugging designated as embryonic day (E) 0.5. PCR genotyping was performed with the following primers: wild-type forward primer: TCT GGC TCA CGA TTC TGA AG; mutant forward primer: AAG GTG GCT ACA CAG GCA AG and common reverse primer: CGG AGG TCA AGC TGT TAA GG.

### Embryo collection

Embryo trunks or postnatal nerves were dissected in ice-cold phosphate-buffered saline (PBS), and then fixed overnight in 4% paraformaldehyde (PFA)/PBS. Fixed tissue was washed in PBS, immersed in 20% sucrose/PBS overnight, and then blocked in O.C.T™ (Tissue-Tek®, Sakura Finetek U.S.A. Inc., Torrance, CA) before storing at −80°C. Blocked tissue was sectioned on a Leica cm3050s cryostat (Richmond Hill, ON) at 10 µm and collected on SuperFrost™ Plus slides (Thermo Scientific).

### Peripheral nerve crush

Peripheral nerve crush injuries were performed on the sciatic nerve of P21 wild-type and *Etv5*^−/−^ mice as previously described (Balakrishnan et al., 2016). Briefly, P21 animals were anesthetized (5% isofluorane for induction and 2% for maintenance), hindlimbs were shaved and sterilized, and a small incision was made to expose the sciatic nerve. A crush injury was performed using #10 forceps for one minute, and then the muscle and skin were sutured back together. Buprenorpine subcutaneous injection of 0.1 mL (100 μL of 0.03 mg/mL) were administered for pain on the day of surgery and for 4 days following, with the nerve harvested on day 5.

### Immunohistochemistry

Sections were thawed, rinsed in PBS to remove excess O.C.T., permeabilized in PBT (PBS with 0.1% TritonX), and then blocked in 10% normal horse serum/PBT for 1 hour. Primary antibodies were then diluted in blocking solution and incubated on sections overnight at 4°C, followed by three PBT washes. Species-specific secondary antibodies, conjugated to Alexa 488 or Alexa 555, were diluted 1/500 in PBT and applied to sections for 1 hour. Sections were washed three times in PBT and stained with 4′,6-diamidino-2-phenylindole (DAPI; Santa Cruz Biotechnology) (1:5000 in PBT). Sections were washed three times in PBS and mounted in AquaPolymount (Polysciences). Primary antibodies included: rabbit anti-Etv5 (Abcam ab102010; 1:300), rabbit anti-Tfap2a (Abcam; ab52222; 1:200), goat anti-Oct6 C-20 (Santa Cruz Biotechnology; sc-11661; 1:50), rabbit anti-Sox9 (Millipore; AB5535; 1:500), goat anti-Sox10 (Santa Cruz Biotechnology; sc-17343; 1:400), rabbit anti-Sox10 (Abcam; AB227680; 1:200), Gfap (Dako Cytomation; #Z0334; 1:500) Ngfr (Millipore; #07-476;1:500); BFABP (Millipore; ABN14; 1:500) and mouse anti-NeuN (Millipore MAB377; 1:200).

### RNA in situ hybridization

A digoxygenin-labeled *Etv5* riboprobe was generated as previously described using a 10× labelling mix and following the manufacturer’s instructions (Roche) (Li et al., 2014). The probe was hybridized overnight, and washing and staining procedures were followed as previously described (Touahri et al., 2015).

### Microscopy and image analysis

Images were captured with a QImaging RETIGA EX digital camera and a Leica DMRXA2 optical microscope using OpenLab5 software (Improvision; Waltham MA). Image processing and analysis was performed using Image J software. Three images per wild-type and *Etv5*^−/−^ embryo/nerve were assessed. DAPI channel images were converted into 8-bit format and the threshold was set using weighted mean intensity. Images were inverted for binary conversion. In E18.5 sections, the dorsal root ganglionic region was manually selected using free-form selection tool, while in nerve sections the entire area was assessed. The number of DAPI^+^ cells in the selected region were calculated using particle analysis option. The colocalization of DAPI with the green channel (Sox10^+^/NeuN^+^ cells) was calculated manually.

### Statistical analysis

A minimum of three biological replicates were carried out for all assays. Statistical analysis and graphs were generated using GraphPad Prism 8 software. Student’s t-test (when comparing two groups) or One-way ANOVA with TUKEY post corrections (when comparing groups of more than two) were used. All data expressed as mean value ± standard error of the mean (S.E.M.). In all experiments, a p value <0.05 was considered statistically significant, *p < 0.05, **p < 0.01, ***p < 0.001, and ****p < 0.0001.

## RESULTS

### Schwann cell precursors develop normally in E12.5 *Etv5*^*−/−*^ peripheral ganglia

Previous reports have documented that *Etv5* is expressed in the Schwann cell lineage from as early as E9.0, first appearing in NCCs, and persisting until E12.5 in satellite glia and Schwann cell precursors (SCPs) in the developing DRG flanking the spinal cord (Fig. 1*A,B*) (Hagedorn et al., 2000; Balakrishnan et al., 2016). We confirmed the expression of *Etv5* in SCPs and satellite glia using RNA in situ hybridization (Fig. 1*C*) and by co-immunolabeling with Sox10 (Fig. 1*D*), which marks the Schwann cell lineage at all developmental stages (Balakrishnan et al., 2016). To next assess whether *Etv5* is required during this early temporal window for Schwann cell lineage development, we examined E12.5 wild-type and *Etv5*^*tm1Kmm*^ mutant embryos (Fig. 1*B*). Notably, we studied *Etv5*^*tm1Kmm*^ mutant mice carrying a deletion of exons 2-5 encoding the initiation codon and a transactivation domain (hereafter *Etv5*^−/−^) because animals homozygous for a null allele (*Etv5*^*tm1Hass*^), which lack the Etv5 DNA binding domain, die by E8.5 (Chen et al., 2005; Kuure et al., 2010), precluding an analysis of Schwann cell lineage development.

**Figure 1.**
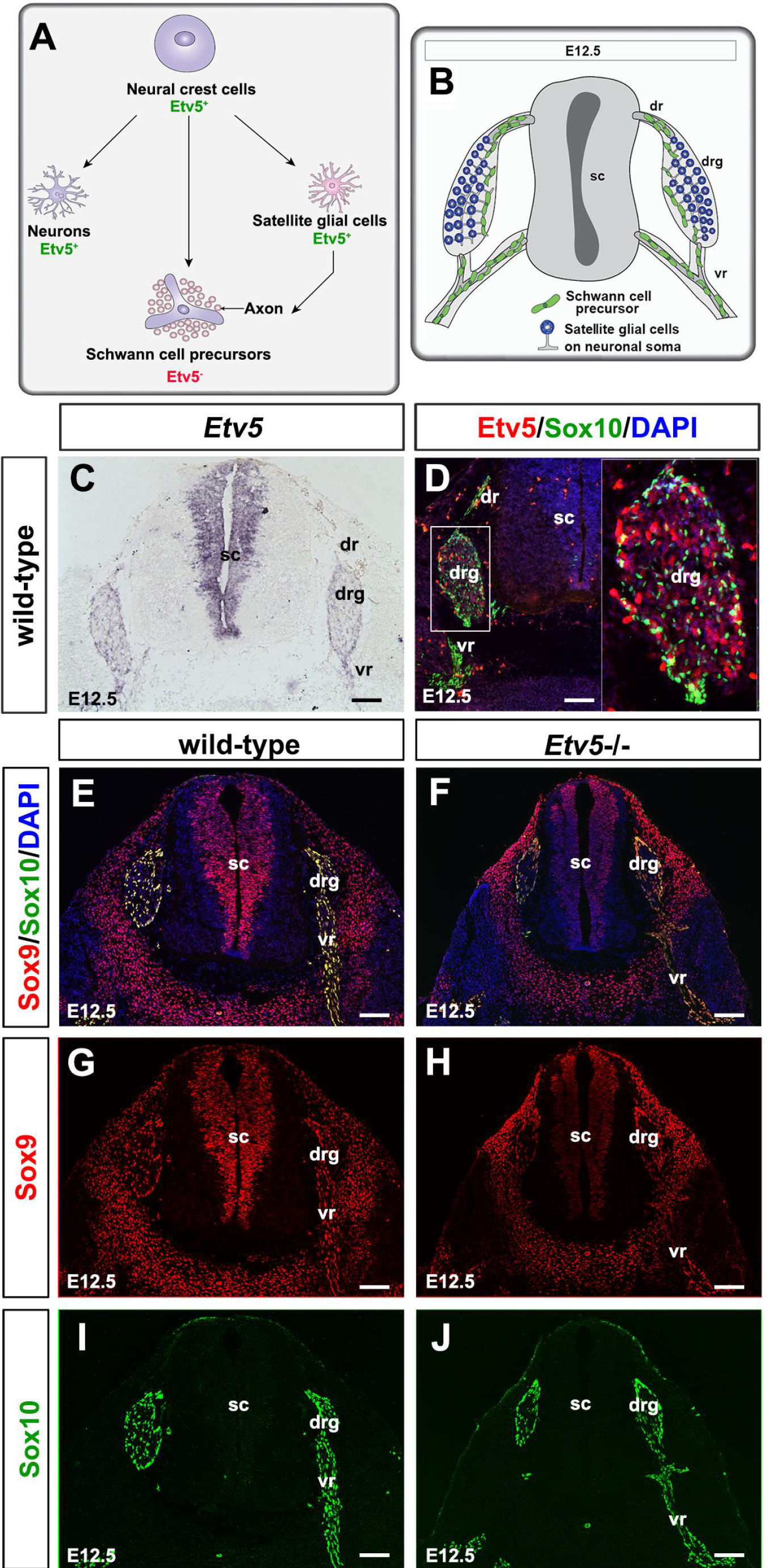
Schwann cell transcriptional regulators are expressed normally in E12.5 *Etv5*^−/−^ Schwann cell precursors (SCPs). Etv5^+^ neural crest cells give rise to Etv5^+^ sensory neurons, Etv5^+^ satellite glia, and Etv5^−^ Schwann cell precursors (A). Schematic representation of E12.5 trunk section (B). Distribution of *Etv5* transcripts in E12.5 transverse sections of the spinal cord (C). Co-expression of Etv5 (red, D), Sox9 (red, E-G,I) and Sox10 (green, D,E,F,H,J) with DAPI counterstain (blue) in E12.5 wild-type (D,E,G,I) and *Etv5*^−/−^ (F,H,J) embryos. Inset in (D) presents 3X magnified image of region marked by a dotted box. dr, dorsal root; drg, dorsal root ganglion; sc, spinal cord; vr, ventral root. Scale bars, 60 μm.

We first examined the expression of Sox9 and Sox10, two SRY-box family HMG transcription factors. Sox10 is a specific marker of the Schwann cell lineage, but also marks early migrating NCCs and satellite glia throughout the embryonic period and into postnatal stages and is required for development past the immature Schwann cell (iSC) stage (Kuhlbrodt et al., 1998; Finzsch et al., 2010; Balakrishnan et al., 2016). Sox9 is also expressed early on in the Schwann cell lineage, beginning at the NCC stage (Cheung and Briscoe, 2003; Balakrishnan et al., 2016), and it regulates NCC development (Cheung and Briscoe, 2003). In E12.5 wild-type embryos, Sox9 was expressed more widely, marking SCPs coalescing in the developing DRG and ventral root, but also labelling neural progenitors in the neural tube, part of the CNS, as well as other migratory NCC populations migrating over the surface ectoderm, and mesenchymal cells (Fig. 1 *E,G*). A very similar pattern of Sox9 expression was observed in E12.5 *Etv5*^−/−^ embryos, including in SCPs in the DRG, ventral root, and the motor root of the spinal nerve (Fig. 1*F,H*). In E12.5 sections through the trunk, Sox9 expression overlapped with Sox10 in the DRG and ventral root, but Sox10 expression was much more restricted to the Schwann cell lineage (Fig. 1*E,I*). A similar pattern of Sox10 expression was seen in E12.5 *Etv5*^−/−^ embryos, with Sox10^+^ cells restricted to glial cells in the DRG and ventral root (Fig. 1*F,J*).

Transcription factors initiate developmental programs by turning on the expression of genes with functional roles in fate specification and differentiation. Brain fatty acid binding protein (BFABP) is the earliest ‘glial’ gene turned on in SCPs in a Sox10-dependent manner (Finzsch et al., 2010). Another early marker is Ngfr (also known as p75NTR), a common receptor for neurotrophins, the deletion of which results in a reduced size of peripheral ganglia and reduction in Schwann cell number (von Schack et al., 2001). In E12.5 wild-type embryos, both BFABP (Fig. 2*A,A*’) and Ngfr (Fig. 2*C,C*’) were co-expressed with Sox10 in SCPs in the DRG and also prominently in the dorsal root, the sensory root of the spinal nerve. BFABP also marked the dorsal neural tube and floor plate, while Ngfr was detected more prominently in the ventral neural tube. Very similar patterns of expression were observed in E12.5 *Etv5*^−/−^ embryos, including prominent expression of BFABP (Fig. 2*B,B*’) and Ngfr (Fig. 2*D,D*’) in Sox10^+^ SCPs in the DRG and dorsal root.

**Figure 2.**
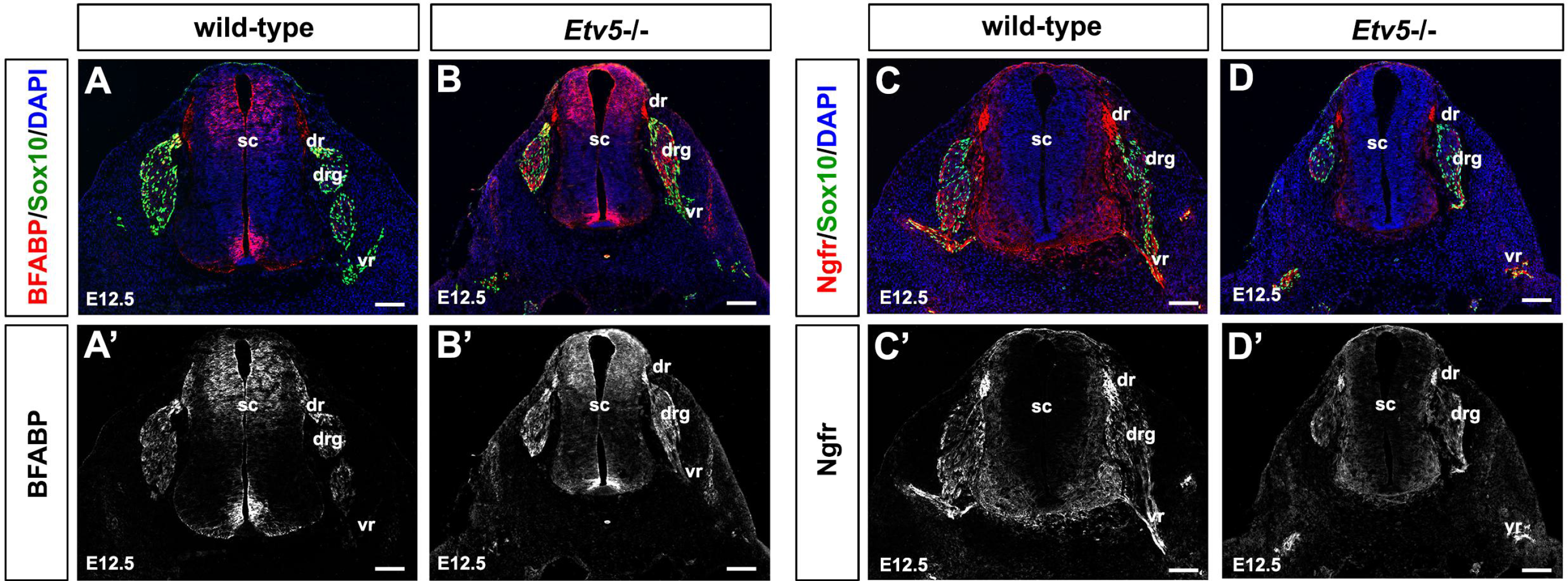
Schwann cell non-transcription factor lineage markers are expressed normally in E12.5 *Etv5*^−/−^ SCPs. Co-expression of Sox10 (green, A-D) with BFABP (red, A,B, black/white, A’,B’) and Ngfr (red, C,D, black/white, C’,D’), counterstained with DAPI (blue, A-D) in E12.5 wild-type (A,A’,C,C’) and *Etv5*^−/−^ (B,B’,D,D’) transverse sections through the trunk. dr, dorsal root; drg, dorsal root ganglion; sc, spinal cord; vr, ventral root. Scale bars, 60 μm.

Thus, at E12.5, there are no notable defects in the general positioning or SCP-specific gene expression in *Etv5*^−/−^ mutant DRG, ventral, and dorsal roots.

### Immature Schwann cells develop normally in E15.5 *Etv5*^−/−^ peripheral ganglia

Between E12.5 and E15.5, some SCPs persist while others proceed on to form immature Schwann cells (iSCs) that populate the developing spinal ganglia and nerves (Jessen and Mirsky, 2005). We focused on the lumbar spinal cord in E15.5 wild-type and *Etv5*^*−/−*^ embryos (Fig. 3*A,B*), and again examined the expression of Sox9 (Fig. 3*C,D*) and Sox10 (Fig. 3*E,F*). Both Sox9 and Sox10 were expressed in a characteristic salt-and-pepper expression pattern in scattered SCPs and iSCs throughout the DRG and in the dorsal and ventral roots in E15.5 wild-type and *Etv5*^*−/−*^ embryos (Fig. 3*C-F*). We also examined the expression of two additional transcription factors with a later onset of expression in SCPs, including transcription factor AP-2α (Tfap2a) (Stewart et al., 2001; Balakrishnan et al., 2016) and Sox2, an inhibitor of Schwann cell myelination that is expressed in SCPs and iSCs, but not in pro-myelinating and myelinating Schwann cells (Le et al., 2005; Adameyko et al., 2012; Balakrishnan et al., 2016). Tfap2a was expressed throughout the E15.5 wild-type DRG (Fig. 3*G*), while Sox2 was only detected in a few Sox10^*+*^ SCPs and iSCs (Fig. 3*I,I’*), contrasting to the robust expression of Sox2 in the spinal cord ventricular zone (Fig. 3*I,I’*). Notably, both Tfap2a (Fig. 3*H*) and Sox2 were similarly expressed in the Schwann cell lineage in E15.5 *Etv5*^−/−^ DRGs (Fig. 3*J,J’*).

**Figure 3.**
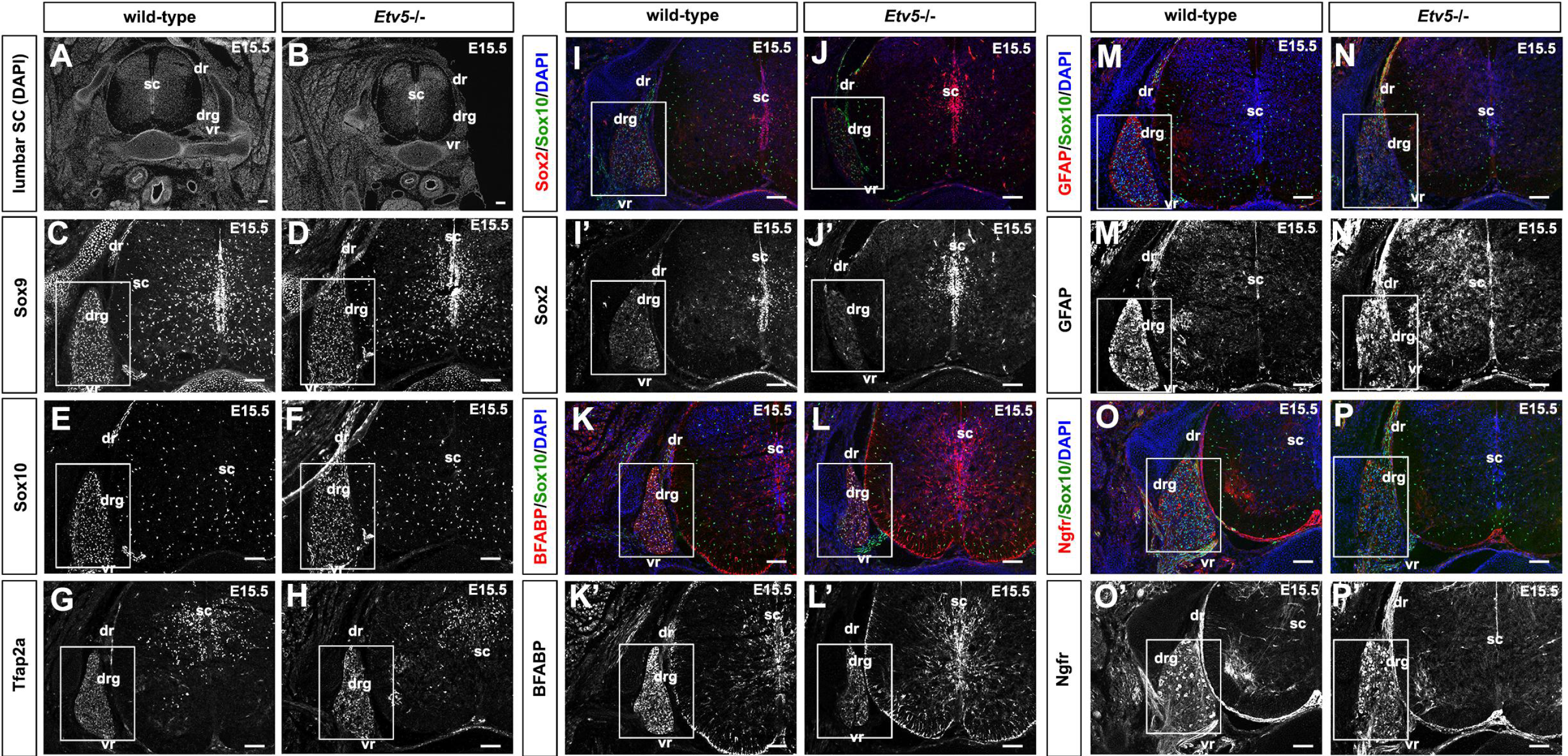
Schwan cell lineage markers are expressed normally in E15.5 *Etv5*^−/−^ immature Schwann cells (iSCs). Low magnification DAPI-stained images of transverse sections through the lumbar spinal cord of E15.5 wild-type (A) and *Etv5*^−/−^ (B) embryos. (C-H) Expression of Sox9, Sox10 and Tfap2a in E15.5 wild-type (C,E,G) and *Etv5*^−/−^ (D,F,H) transverse sections through the lumbar spinal cord. (I-P) Co-expression of Sox10 (green, I-P) with Sox2 (red, I,J, black/white, I’,J’), BFABP (red, K,L, black/white, K’,L’), GFAP (red, M,N, black/white, M’,N’), and Ngfr (red, O,P, black/white, O’,P’), counterstained with DAPI (blue, I-P) in E12.5 wild-type (I,I’,K,K’,M,M’,O,O’) and *Etv5*^−/−^ (J,J’,L,L’,N,N’,P,P’) transverse sections through the lumbar spinal cord. dr, dorsal root; drg, dorsal root ganglion; sc, spinal cord; vr, ventral root. Scale bars (A,B), 100 µm; (C-P’), 60 μm.

Finally, to assess the maturation process of SCPs into iSCs, we examined the expression of non-transcriptional regulators that mark the Schwann cell lineage, including BFABP (Fig. 3*K,L,K’,L’*), the intermediate filament, glial fibrillary acidic protein (GFAP) (Fig. 3*M,M’,N,N’*) and Ngfr (Fig. 3*O,O’,P,P*’). All three of these proteins were expressed in scattered Sox10^+^ SCPs and iSCs in both E15.5 wild-type (Fig. 3*K,K’,M,M’,O,O*’) and *Etv5*^*−/−*^ (Fig. 3*L,L’,N,N’,P,P*’) DRGs. Thus, SCPs and iSCs develop normally in *Etv5* hypomorphic mutants.

### Late immature Schwann cells/pro-myelinating Schwann cells are detected in E18.5 *Etv5*^−/−^ peripheral ganglia and in the dorsal and ventral roots

By E18.5, some iSCs persist, while other iSCs begin to associate with large-diameter axons to become pro-myelinating Schwann cells, whereas iSCs in contact with smaller diameter axons become non-myelinating Schwann cells (Feltri et al., 2015). We first labelled all Schwann cells in the lineage with Sox10 in E18.5 wild-type (Fig. 4*A*) and *Etv5*^*−/−*^ (Fig. 4*B*) transverse sections through the lumbar spinal cord, and detected similar numbers of Sox10+ Schwann cells in in the DRG, as well as in the dorsal and ventral roots in the wild-type and *Etv5*^*−/−*^ mutants (Fig. 4*I*). Further, we examined whether the neuronal cells populating the DRG were affected by labelling the cells with NeuN, a neuronal marker. A similar number of NeuN^+^ neurons were observed in both the wild-type and *Etv5*^*−/−*^ mutant DRGs (Fig. 4*C,D,J*). Next, we examined the expression of Pou3f1 (Oct-6), which is a marker of late iSCs/pro-myelinating Schwann cells that is required for the transition to a myelinating phenotype transition (Arroyo et al., 1998). We detected Pou3f1 expression in the ventral roots in both E18.5 wild-type (Fig. 4*E*) and *Etv5*^*−/−*^ (Fig. 4*F*) embryos, suggesting that Sox10 cells mature to a myelinating stage. Finally, to detect satellite glial cells in the developing DRG, we examined the expression of glutamine synthetase (Ohara et al., 2009), revealing that this marker was expressed in both wild-type (Fig. 4*G*) and *Etv5*^*−/−*^ (Fig. 4*H*) mutant DRGs. Thus, in *Etv5* hypomorphic mutants, normal numbers of Schwann cells and DRG neurons are generated during development.

**Figure 4.**
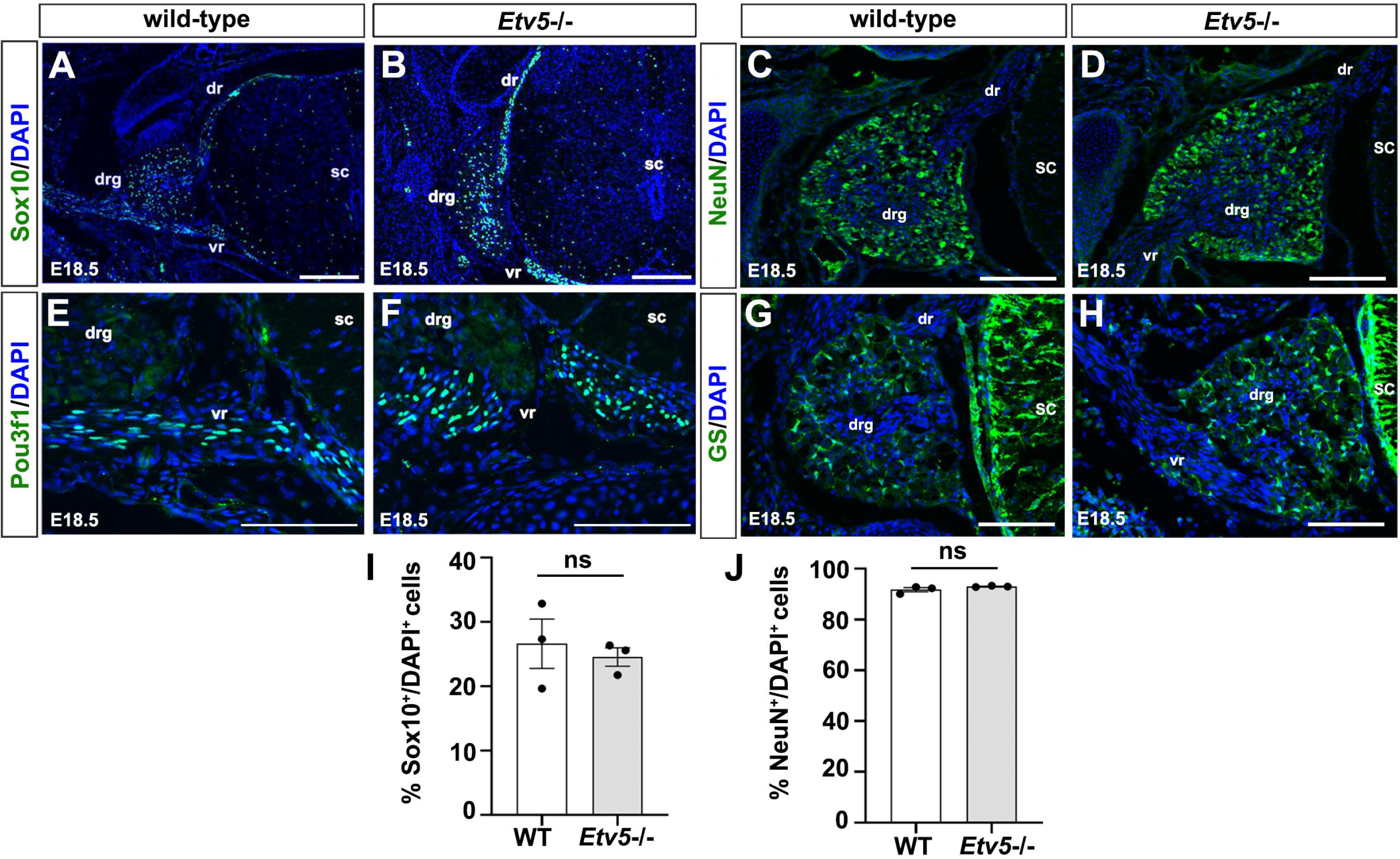
Schwan cell lineage markers are expressed normally in E18.5 *Etv5*^−/−^ late immature/pro-myelinating Schwann cells. Labelling of Sox10 (A,B), NeuN (C,D), Pou3f1 (E,F), and Glutamine Synthetase (GS; G,H) with DAPI counterstain in E18.5 wild-type (A,C,E,G) and *Etv5*^−/−^ (B,D.F.H) transverse sections through the lumbar spinal cord. Quantification of percentage of Sox10^+^/DAPI^+^ cells (I) in the dorsal root ganglia, dorsal and ventral root, and NeuN^+^/DAPI^+^ cells (J) in the dorsal root ganglia. Error bars=s.e.m.. drg, dorsal root ganglion; sc, spinal cord; vr, ventral root; ns, non-significant. Scale bars, (A,B), 60 μm; (C,D,G.H), 40 µm; (E,F), 20 µm.

### Schwann cells populate the postnatal sciatic nerve in *Etv5*^−/−^ mice and respond to injury with an expansion in number

As a final question, we asked whether there were defects in Schwann cells found in the early postnatal nerve, at postnatal day (P) 21, when most pro-myelinating Schwann cells have converted to a myelinating phenotype (Balakrishnan et al., 2016). At P21, we observed expression of Sox10 in scattered cells throughout longitudinal sections of the sciatic nerve in both wild-type (Fig. 5*A*) and *Etv5*^*−/−*^ (Fig. 5*B*) animals, and there were no differences in Schwann cell numbers between these groups in the uninjured nerve (Fig. 5*E*). We then subjected both P21 wild-type and *Etv5*^*−/−*^ animals to a sciatic nerve crush injury, which induces a de-differentiation and subsequent expansion of Schwann cells in the distal stump by 5 days post-injury (dpi) (Balakrishnan et al., 2016). We observed an increase in the number of Sox10^+^ Schwann cells in both P21 wild-type (p=0.028; Fig. 5*C,E*) and *Etv5*^−/−^ (p=0.0001; Fig. 5*D,E*) distal stumps at 5 dpi, suggesting that the repair response occurs normally in *Etv5*^−/−^ nerves. Interestingly, a significant increase in Sox10^+^ cells was observed in *Etv5*^−/−^ injured nerves compared to wild-type injured nerves (p=0.0027; Fig. 5*E*), indicating that *Etv5* normally inhibits the ability of Schwann cells to expand post injury.

**Figure 5.**
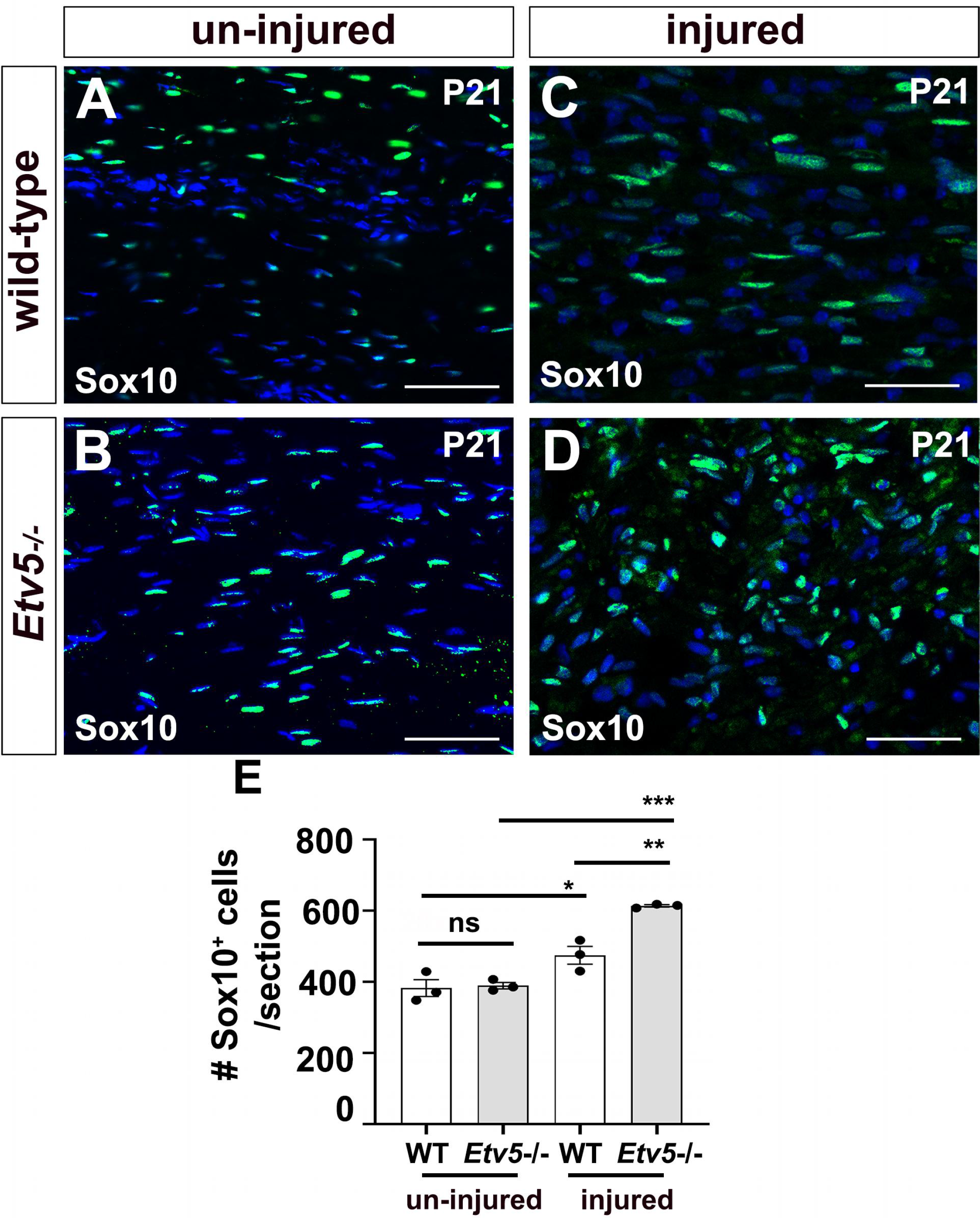
Increase in Sox10^+^ Schwann cells post-injury in *Etv5*^−/−^ sciatic nerve. Sox10 expression in longitudinal sections of the uninjured (A,B) and injured (C,D) P21 sciatic nerve from wild-type (A,C) and *Etv5*^−/−^ (B,D) animals. Quantification of number of Sox10^+^ cells (E) in the entire section. Error bars=s.e.m. *, p < 0.05; **, p < 0.01; ***, p < 0.005. Scale bars, 20 μm.

Taken together, these studies suggest that while the Schwann cell lineage develops normally in *Etv5*^*−/−*^ embryos, from embryonic to early postnatal stages, the response of *Etv5*^*−/−*^ Schwann cells to injury is altered, with an increase in Sox10^+^ Schwann cells populating near the nerve injury site.

## DISCUSSION

In this study we investigated the role of *Etv5* in Schwann cell development using a genetic mutant that deletes exons 2-5, an allele associated with defects in stem cell self-renewal in the spermatogonial lineage (Chen et al., 2005). Using a panel of well annotated Schwann cell markers and quantitative studies at E18.5, we did not observe any notable defects in Schwann cell or neuronal differentiation in *Etv5*^*−/−*^ peripheral ganglia at embryonic stages. However, following an acute peripheral nerve injury, which results in the de-differentiation of mature Schwann cells and an increase in Schwann cell proliferation (Jessen and Mirsky, 2019b), more Sox10^+^ cells were observed in *Etv5*^*−/−*^ nerves, implicating *Etv5* as a negative regulator of the Schwann cell injury response. ERK1/2 signal transduction is activated downstream of NRG1 (Grossmann et al., 2009; Sheean et al., 2014), which is a critical regulator of myelination, and ERK1/2 induce Schwann cell myelination (Ishii et al., 2013; Sheean et al., 2014), including after injury (Harrisingh et al., 2004; Guertin et al., 2005; Napoli et al., 2012; Kim et al., 2013). Our findings were therefore surprising as our expectation was that *Etv5* would be a positive regulator of the injury response.

Notably, in our study we used an *Etv5* mutant allele that has a deletion of exons 2-5 (*Etv5*^*tm1Kmm*^), which results in a very striking reduction in spermatogonial stem cell self-renewal (Chen et al., 2005). In contrast, an *Etv5* mutation that removes the DNA binding domain (*Etv5*^*tm1Hass*^) is embryonic lethal at E8.5, suggesting it is a true null allele (Chen et al., 2005), but precluding us from analysing Schwann cells due to the early embryonic lethality. Due to the embryonic lethality associated with ‘more severe’ mutant alleles of *Etv5*, it is believed that our studied *Etv5* allele (exon 2-5 knocked out) is hypomorphic rather than a true null allele. Further support for the designation of *Etv5*^*tm1Kmm*^ as a hypomorphic allele comes from the lack of a developmental kidney defect (Lu et al., 2009), whereas the generation of chimeric embryos using *Etv5*^*tm1Hass*^ embryonic stem cells revealed that *Etv5* is required for kidney development (Kuure et al., 2010). Future studies on *Etv5* function in the Schwann cell lineage would require either the use of such a chimeric approach, or a genetic approach, such as the use of a floxed allele of *Etv5* (Zhang et al., 2009). However, even using these alternative approaches, we may not observe a Schwann cell developmental phenotype if there is genetic redundancy. Indeed, different members of the Fgf-synexpression group of ets transcription factors (*Etv1, Etv4, Etv5*) may compensate for one another to some extent, as shown for *Etv4* and *Etv5* in kidney development (Kuure et al., 2010). Thus, we may have not uncovered a role for *Etv5* in developing Schwann cells due to issues of genetic redundancy. Nevertheless, despite these caveats, we can conclude that the reduced levels of *Etv5* function associated with the *Etv5*^*tm1Kmm*^ allele, which has striking phenotypic consequences in other lineages, does not impact early Schwann cell development, but does impact the Schwann cell response to injury.

An important area for future studies will be to further examine the role of *Etv5* in mature Schwann cells. Schwann cell transplants have the potential to aid peripheral nerve repair, and efforts are being made to improve the isolation and expansion of these cells to provide an adequate source for repair purposes. In this regard it is interesting that both human nerve-derived and skin derived Schwann cells cultured *in vitro* express a large number of Schwann cell markers associated with an early developmental phenotype, which includes *Etv5* (Stratton et al., 2017). In our study here we provide the first glimpse that a decline in *Etv5* expression leads to more number of Schwann cells following a peripheral nerve crush injury. One possibility is that the knockdown of this factor could thus be exploited for regenerative purposes.

RTK-ERK signaling is crucial for Schwann cell differentiation (Newbern et al., 2011) and has been implicated in the myelination process, in part by inducing the expression of pro-myelinating transcription factors such as *Yy1* (He et al., 2010). Other ets-domain transcription factors that are involved in ERK signaling are *Etv1* and *Etv4*. While *Etv1* is expressed in myelinating Schwann cells (Srinivasan et al., 2007), it is not required for Schwann cell myelination (Fleming et al., 2016). In this regard, it is interesting to note that dominant negative *Etv5* misexpression in NCC cultures impacts neuronal fate specification, whereas glial fates are left unperturbed (Paratore et al., 2002). Our study similarly indicated that *Etv5* is not required for generation of mature Schwann cells. In contrast, the increase in Sox10^+^ cell population in *Etv5*^*−/−*^ sciatic nerve post-injury (compared to injured wild-type nerves) suggested that loss of *Etv5* expression may promote Schwann cell de-differentiation post-injury. Since ERK1/2 signaling is involved in both Schwann cell myelination (Ishii et al., 2013) as well as in promoting a de-differentiated Schwann cell state (Kim et al., 2013), the exact role of Etv5 needs to be further investigated. Questions to be addressed in the future include whether Schwann cells myelinate axons normally in *Etv5*^*−/−*^ animals, and if *Etv5* regulates Schwann cell proliferation post-injury in conjunction with *Sox2* and *Jun* activity, which play an important role in promoting the de-differentiated repair Schwann cell phenotype (Jessen and Mirsky, 2019b).

In summary, while our study does not support a critical role for *Etv5* in Schwann cell development, we demonstrate that *Etv5* is involved in regulating the repair Schwann cell response post-injury.

## Acknowledgments

C.S. holds the Dixon Family Chair in Ophthalmology Research at the Sunnybrook Research Institute. L.B. is funded by a CIHR *Fredrick Banting and Charles Best graduate award – Master’s Program*.

